# *Rickettsia typhi* peptidoglycan mapping with data-dependent tandem mass spectrometry

**DOI:** 10.1101/2022.03.10.483817

**Authors:** Benjamin L. Oyler, Kristen E. Rennoll-Bankert, Victoria I. Verhoeve, M. Sayeedur Rahman, Abdu F. Azad, Joseph J. Gillespie, David R. Goodlett

## Abstract

*Rickettsia* species are diverse Gram-negative obligate intracellular bacteria often pervasive in numerous invertebrates, as well as fungal, nematode and microeukaryotic hosts. Certain species are etiological agents for well-known arthropod-borne illnesses; e.g., *R. rickettsii* (Rocky Mountain Spotted Fever), *R. prowazekii* (Epidemic Typhus), and *R. typhi*, (Endemic Typhus). Living freely in eukaryotic cytosol presumably exposes rickettsiae to host cell immune receptors, particularly those recognizing bacterial cell envelope glycoconjugates. However, the mechanics of host recognition of rickettsiae remain poorly defined. As rickettsiae synthesize a canonical Gram-negative cell envelope that includes peptidoglycan (PGN) and lipopolysaccharide (LPS), structural insight on these macromolecules is important for deciphering host responses to these pathogens. In this work, PGN from *R. typhi* was digested and the resultant subunits were analyzed by two different, albeit complementary, sample preparation methods. Both approaches were subsequently subjected to liquid chromatography/mass spectrometry analysis to infer PGN structure. *R. typhi* PGN was determined to be similar to most other Gram-negative bacteria, with mDAP-type muropeptide subunits. However, additional alanine residues were observed elongating the muropeptide stems, rather than the glycine residues usually observed in Gram-negative bacterial PGN. Despite this deviation, *R. typhi* contains a murein layer that is predicted to agonize host cellular PGN receptors and be susceptible to PGN-targeting antimicrobials. This same structure is likely synthesized by all *Rickettsia* species, as bioinformatics and comparative genomics analyses indicate the biosynthesis of PGN is highly conserved. Determining how host cells process this canonical glycoconjugate during infection is crucial for identifying factors behind rickettsial pathogenesis, including immunoavoidance or proinflammatory mechanisms possibly employed by rickettsiae with varying pathogenic potential.

## Introduction

*Rickettsia typhi* (Alphaproteobacteria; Rickettsiales) is an obligate intracellular bacterial pathogen of mammals and is responsible for murine (or endemic) typhus. *R. typhi* and the closely related agent of epidemic typhus, *R. prowazekii*, comprise the **Typhus Group** (**TG**) rickettsiae and are found in blood-feeding species of fleas and lice (Azad et al., 1997) *R. typhi* is mostly transmitted to humans through the feces of infected fleas (Azad and Beard, 1998). Clinical diagnosis of murine typhus is difficult because its symptoms are not specific and often misdiagnosed by physicians in the United States due to its relatively low incidence in urban areas (Civen and Ngo, 2008). Symptoms of murine typhus usually present 7-14 days after infection and include fever, headache, rash, and myalgia (Osterloh, 2017). As with other Rickettsioses, doxycycline is the treatment of choice although its effect weakens later in infection. Effective vaccines do not exist (Osterloh, 2020).

Rickettsiae are small and rod-shaped, with a typical Gram-negative cell envelope structure and a large electron-translucent S-layer (Silverman and Wisseman, 1978). Early biochemical studies revealed a lack of glycolytic activity (Coolbaugh et al., 1976), yet paradoxically, the presence of certain components of **peptidoglycan** (**PGN**) and **lipopolysaccharide** (**LPS**) in the *Rickettsia* cell envelope (Amano et al., 1993; Pang and Winkler, 1994). More recently, *Rickettsia* comparative genomics analyses revealed the presence of complete pathways for PGN and LPS biosynthesis (Gillespie et al., 2008, 2012b), leading to the hypothesis that rickettsiae pilfer host amino sugars to initiate biosynthesis of these macromolecules (Driscoll et al., 2017). Despite *R. typhi* (McLeod et al., 2004) and *R. prowazekii* (Andersson et al., 1998) genomes carrying far less genes than other rickettsiae (Gillespie et al., 2008, 2015), their PGN and LPS synthesis pathways are highly conserved.

The primary sites of infection with *R. typhi* are usually vascular endothelial cells near to a flea-bite locus. *R. typhi* readily invades endothelial cells to grow and proliferate until causing host cell lysis (Sahni and Rydkina, 2009), after which bacteria travel throughout the circulation and infect almost all host cell types (Osterloh, 2017). Curiously, *R. typhi* infects macrophages (MΦ) and dendritic cells, first-line responders of the innate immune system (Radulovic et al., 2002). In particular, MΦ are central for either clearance of the infection or succumbing to pathogen colonization, which results in systemic bacterial spread (Sahni et al., 2019). Since *R. typhi* displays several known innate immune **pattern recognition receptor** (**PRR**) ligands (i.e. PGN and LPS) in its cell envelope, it is phagocytosed by possible recognition of these pathogen markers, after which it is able to evade the killing mechanisms of the macrophage.

PGN is a polymer of disaccharide β-linked ***N*-acetylglucosamine** (***N*AG**) and ***N*-acetylmuramic acid** (***N*AM**) subunits covalently bound to a short “stem” peptide at the C-3 position of *N*AM and cross-linked with short peptides. This structure comprises a thin layer in the periplasm between the inner and outer Gram-negative bacterial membranes (often referred to as the murein layer) and forms a thick layer on the outer leaflet of Gram-positive bacterial membranes between the cell and extracellular milieu (Schleifer and Kandler, 1972). Thus, Gram-positive bacteria are defined by their ability to retain crystal violet stain in their thick PGN layer. The peptide cross-links in Gram-negative bacteria, like *R. typhi*, usually consist of a direct link between one D-alanine residue and one **meso-diaminopimelic acid** (**mDAP**) residue on two distinct PGN stem peptides (Vollmer et al., 2008). However, there is structural variability in both *N*AG-*N*AM modification and variance in stem peptide composition in both Gram-negative and Gram-positive bacteria (Park and Uehara, 2008), warranting the direct determination of PGN structures over genome sequence-based predictions for PGN biosynthesis pathways.

PGN subunits, namely ***N*-acetyl-d-glucosaminyl-*N*-acetyl-d-muramyl-l-alanyl-d-isoglutamyl-meso-diaminopimelic acid** (**GM-TriDAP**, Gram-negative bacteria) and **muramyl dipeptide** (**MDP**, Gram-positive and Gram-negative bacteria), are ligands for such PRRs as **nucleotide-binding oligomerization domain-containing proteins 1 and 2** (**NOD1** and **NOD2**, respectively) (Girardin et al., 2003a, 2003b), as well as other **PGN recognition proteins** (**PGRPs**). NOD1 and NOD2 are two members of a larger family of PRRs called **NOD-like receptors** (**NLRs**), that are responsible for acute phase inflammation events following bacterial infection. NOD1 or NOD2 activation by PGN results in NF-κB translocation to the nucleus where it facilitates expression and release of cytokines and chemokines (McCarthy et al., 1998; Bertin et al., 2000). NLRs act in concert with other PRRs, such as **Toll-like receptors** (**TLR**), to recognize and clear bacterial infection in the acute phase.

As PGN is absent in most eukaryotes yet found in most bacteria, many successful antibiotics have targeted PGN; however, resistance to PGN-targeting antibiotics is on the rise (Magiorakos et al., 2012; Glen and Lamont, 2021). Structural information is needed to determine if pathogens modify PGN to avoid activating host detection and/or guard against PGN-targeting toxins and antibiotics. Perhaps due to the low yield of PGN from many Gram-negative bacteria, as well as the difficulty of harvesting enough murein layer from rickettsiae grown in host cell culture, PGN structure has not been determined for any *Rickettsia* species. In this work, we report the PGN subunit structure for *R. typhi* to provide concrete molecular data for future host infection immunology studies.

## Materials and Methods

### *R. typhi* culture and PGN isolation

*R. typhi* cells were cultured according to a previously published method (Rennoll-Bankert et al., 2016). Vero76 cells (African green monkey kidney, ATCC: CRL-1587) and HeLa (ATCC: CCL-2) were maintained in minimal Dulbecco’s Modified Eagle’s Medium (DMEM with 4.5 gram/liter glucose and 480 L-glutamine; Mediatech, Inc.), with 10% heat inactivated fetal bovine serum (FBS) added, at 37°C with 5% CO2. Propagation of *R. typhi* strain Wilmington (ATCC: VR-144) was performed in Vero76 cells cultured in DMEM, with 5% heat inactivated fetal bovine serum added, at 34 °C with 5% CO2. Rickettsiae were isolated as described elsewhere (Kaur et al., 2012). *R. typhi* was used at a multiplicity of infection (MOI) of ~100:1 to infect host cells. Prior to host cell infection, cells were washed with DMEM and 5% FBS.

PGN was isolated from *R. typhi* cells using two different established methods (Tong et al., 1997; Kühner et al., 2014). Briefly, the first method (*Experiment 1*) consisted of cell wall isolation by boiling, centrifugation, resuspension in water, sonication, and digestion of residual proteins (Kühner et al., 2014). The second method (*Experiment 2*) utilized a BeadBeater (BioSpec Products, Bartlesville, OK, USA) to isolate bacterial cell wall material with several filtration steps performed afterward. These were followed by resuspension in SDS solution, sedimentation by ultracentrifugation, washing, digestion of residual proteins, and re-sedimentation by ultracentrifugation (Tong et al., 1997).

### PGN digestion and preparation for LC/MS analysis

PGN was also digested using two different established approaches (Billot-Klein et al., 1996; Kühner et al., 2014). The first method (*Experiment 1*) employed only mutanolysin for digestion in a 1/10 volumetric ratio to the sample, followed by reduction of *N*AM and pH adjustment to pH 2-3 (Kühner et al., 2014). The second method (*Experiment 2*) utilized both mutanolysin (250 μg mL-1) and lysozyme (200 μg mL-1) to digest PGN overnight (16 hr), followed by centrifugation (Billot-Klein et al., 1996). Published procedures were strictly followed, except that muropeptide reduction was not performed before LC/MS analysis.

### Data-dependent acquisition of tandem mass spectra

For *Experiment 1*, PGN digests were diluted 100-fold in water and analyzed by LC/MS with a nanoAcquity binary nano-flow LC system (Waters Corporation, Milford, MA, USA) equipped with an autosampler, coupled to an Orbitrap Fusion Tribrid mass spectrometer (Thermo Scientific, San Jose, CA, USA) operated in positive ionization mode. For *Experiment 2*, PGN digests were diluted 100-fold in water and analyzed by LC/MS with an easyNLC 1000 binary nano-flow LC system (Thermo Scientific, San Jose, CA, USA) equipped with an autosampler, coupled to an Orbitrap Fusion Tribrid mass spectrometer (Thermo Scientific, San Jose, CA, USA) operated in positive ionization mode. For both experiments, muropeptide separation was achieved on an in-house fabricated laser pulled tip column consisting of a 100 μm ID x 150 mm length of fused silica (Polymicro Technologies, Phoenix, AZ, USA) packed with YMC Triart (YMC America, Inc., Allentown, PA, USA) 120 Å pore size, C18, 3 μm particles (*Experiment 1*) or 1.9 μm particles (*Experiment 2*). A binary gradient elution program was performed using Optima (Fisher Scientific, Waltham, MA, USA) LC/MS grade water with 0.1% formic acid (Solvent A) and Optima (Fisher Scientific, Waltham, MA, USA) LC/MS grade acetonitrile with 0.1% formic acid (Solvent B) either from 5% Solvent B to 35 % Solvent B in 60 min (*Experiment 1*) or from 5% Solvent B to 35% Solvent B in 30 min (*Experiment 2*). For *Experiment 1*, nominal mass accuracy tandem mass spectra were acquired in **data dependent acquisition mode** (**DDA**) using **collision induced dissociation** (**CID**) at **normalized collision energy** (**NCE**) of 35%. For *Experiment 2*, accurate tandem mass spectra were collected in DDA mode using **higher energy collisional dissociation** (**HCD**) at 25% after quadrupole isolation for tandem experiments, with detection in the Orbitrap. Apex detection was employed to obtain tandem mass spectra at the highest intensity region of chromatographic peaks. DDA mode allows the instrument to choose precursor ions based on the data being acquired in real time, so building lists of precursor ions *a priori* is not necessary. The DDA decision tree for both experiments was designed with a 3 s cycle time in Top Speed configuration, using mass resolution settings of 120K (MS) and 60K (accurate MS/MS) or unit resolution (nominal MS/MS). Only 2+ and above charge states were selected for MS/MS.

### Data analysis

Raw data files were processed in XCalibur software version 3.0 (Thermo Scientific, San Jose, CA, USA). Accurate masses for muropeptides and their product ions were extracted and compared to theoretical monoisotopic masses manually for mass accuracy determination. Proposed chemical structures were constructed in ChemSketch version 14.01 (ACD/Labs, Toronto, ON, Canada). When necessary, data were converted to mzML format using the ProteoWizard msconvert utility (Kessner et al., 2008). Automated peak picking was performed using mMass version 5.5 (www.mmass.org) (Strohalm et al., 2008). Data were plotted using QtiPlot version 0.9.8.9 (www.qtiplot.com) and Figures were generated in Inkscape version 0.91 (https://inkscape.org) and when necessary, modified to an acceptable format using GIMP version 2.8.18 (www.gimp.org).

## Results and Discussion

### Qualitative assessment of muropeptide mass chromatograms

*R. typhi* PGN analysis entailed two approaches for digesting PGN extracted from bacteria grown in Vero cell culture with either mutanolysin alone or both mutanolysin and lysozyme with slightly different downstream protocols (see *Experiments 1* and *2* in the **Materials and Methods**). Observed results from both approaches indicate that both *Experiment 1* and *2* were successful for the isolation and digestion of *R. typhi* PGN extracted from cell culture (**Fig. 1**). **Total ion current chromatograms** (**TICs**) for *Experiment 1* (**Fig. 1A**) and *Experiment 2* (**Fig. 1B**) show the trace for the major muropeptides from *R. typhi* PGN, which have retention times between 15 and 30 min (*Experiment 1*) or 5 and 20 min (*Experiment 2*) during reversed phase gradient elution. Thus, extracted *Rickettsia* PGN is robust to two digestion approaches.

**Figure 1.**
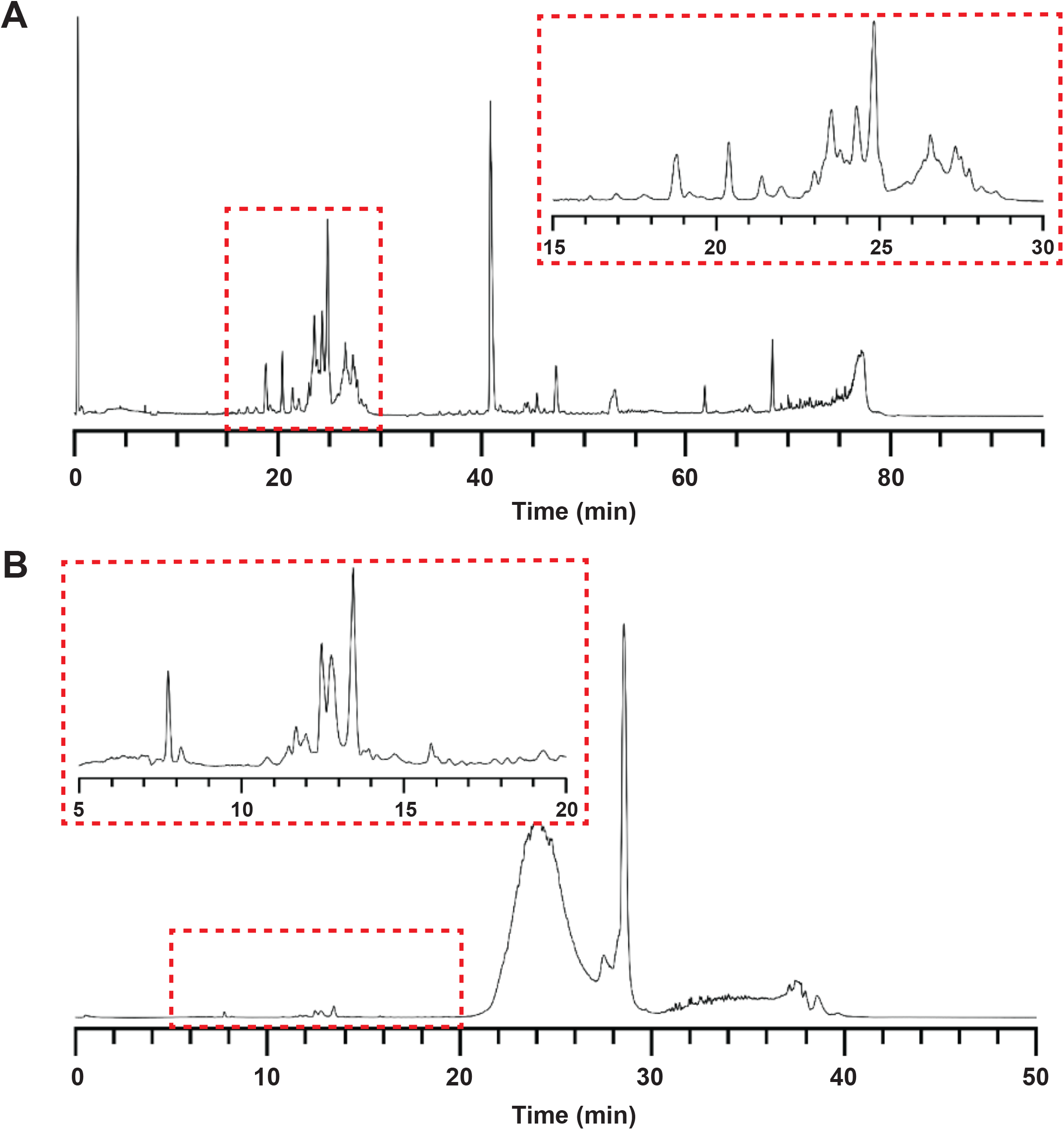
Structure of *R. typhi* PGN derived by two digestion approaches. Total ion current chromatogram (TIC) monitoring ion image current in the Orbitrap mass analyzer for *R. typhi* PGN muropeptides using data dependent acquisition LC-MS/MS from (**A**) *Experiment 1* (mutanolysin alone) or (**B**) *Experiment 2* (mutanolysin and lysozyme). Red dashed boxes: zoomed-in portions of the mass chromatograms showing elution profile of the majority of detected muropeptides. The y-axis represents an arbitrary signal intensity value, so it is not shown.

Since two different sample preparation and chromatographic methods were employed, some comparisons can be made between the data acquired. Although this was not the original purpose of the contrasting approaches, there are some interesting observations that could be useful for future experimental design. First, many of the most abundant peaks in the mass chromatograms are the same; however, there are large differences in both the presence and relative abundance of less abundant peaks. Since the chemistry of both mobile phases and stationary phases remained the same in both experiments, it is unlikely that these observations are due to differences in chromatographic chemistry. Second, and possibly more important, the TIC for *Experiment 2* shows highly abundant, late-eluting peaks which can tentatively be attributed to either intact protein or much larger PGN fragments (**Fig. 1B**). Many charge states of the same molecular weight species were observed in the single stage mass spectra for these peaks (data not shown), indicating numerous basic sites to accept protons when ionized in the positive mode on a mass spectrometer. This observation is common when analyzing ions with many amino acid residues. Third, the separation seems to be faster and possibly more efficient – although it is impossible to make a direct comparison due to sample differences - in *Experiment 2*, in which smaller stationary phase particles were used. This is an expected result for molecules like muropeptides with relatively low molecular weight, but nonetheless notable.

### Evaluation of tandem mass spectra and structure assignment

Tandem mass spectra were acquired for most of the muropeptides in both experiments using DDA-MS approaches. High quality tandem mass spectra were obtained after fine-tuning of the acquisition methods. An exemplar small muropeptide structural assignment was deduced based on both the accurate mass measured in the single stage MS experiment and accurate product ion masses measured in the collisional activation tandem MS experiment (**Fig. 2**). Combined, these data provide high confidence primary structure assignments. As in a prior report (Packiam et al., 2015), collisional activation of short muropeptides produced neutral losses of both hexosamine sugars as well as cleavages of amide bonds as observed with collisional activation of peptides.

**Figure 2.**
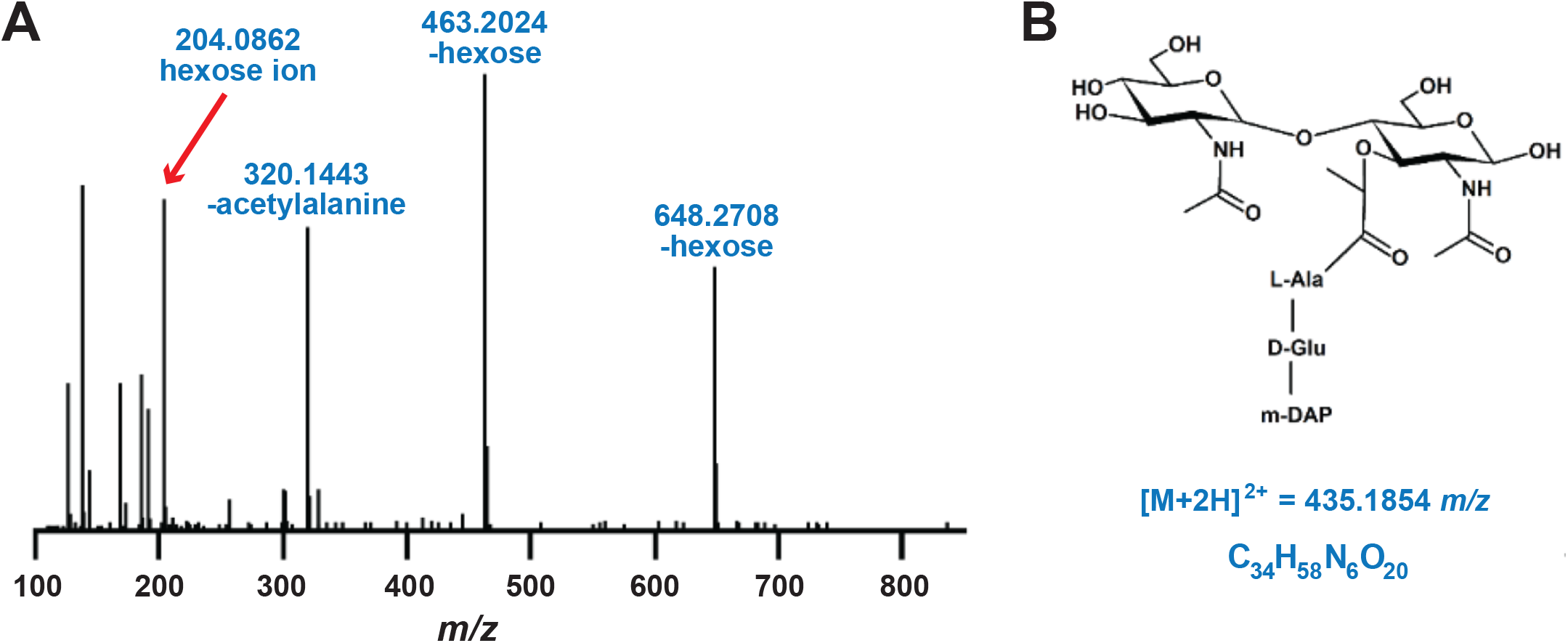
Structure of an *R. typhi* PGN subunit. (**A**) Tandem mass spectrum for a subunit of *R. typhi* PGN consisting of a disaccharide backbone with a tripeptide stem. (**B**) Chemical structure assigned to the precursor ion at m/z 435.1854 from the tandem mass spectrum. The product ion at m/z 463.2024 corresponds to muramyl tripeptide, a good diagnostic ion for mDAP-type muropeptides and Gram-negative bacterial PGN.

Notably, a product ion observed at *m/z* 463.2024 corresponds to a muramyl tripeptide ion, which is a good diagnostic for quickly determining the type of PGN present in the sample. This product ion is only observed in mDAP type PGN – the type produced by most Gram-negative bacteria. With additional method tuning and additional data processing capabilities, most muropeptides containing mDAP would be identifiable by extracting the product ion chromatogram for this ion. In fact, this was the initial approach used in this work to quickly quality control the samples and confirm the presence of PGN fragments (data not shown). It should be noted, however, that optimization of collision energy for different sizes of muropeptides could substantially affect the performance of such a method. More than one LC/MS experiment would probably be needed to cover the entire muropeptide size range. Additionally, an instrument capable of precursor ion scanning, such as a triple quadrupole mass spectrometer, would be able to filter the chromatographic peaks associated with mDAP-type muropeptides during data acquisition.

We were able to confidently predict additional muropeptide structures, for which molecular formulae were assigned based on both single stage and tandem mass spectra (**Table 1** shows an abbreviated list). Since the Orbitrap is an accurate mass analyzer, many chemical formulae could be ruled out using accurate mass measurement alone. From there, it was possible to infer structures based on the tandem mass spectra (**Fig. 2B**). The limitation of this method is that it cannot differentiate between stereoisomers of amino acid residues or sugar residues, so additional chemical analysis would be necessary to derive a muropeptide structure *de novo*. In some cases, additional alanine residues (as in *m/z* 497.2168) were observed attached to the stem peptide, rather than the commonly observed glycine residues. The biological significance of this deviation from typical muropeptide cross-linking is not clear but deserves further investigation.

**Table 1.** Abbreviated list of identified PGN muropeptides with empirical formulae and measured *m/z*.

| Retention time (min) | Precursor charge ($m/z$) | Precursor formula | [M] | $\Delta$ mass (ppm) | Major product ions ($m/z$) |
| --- | --- | --- | --- | --- | --- |
| 7.75 | 435.1854 (+2) | C34H58N6O20 | 868.3552 | 0.23 | 648.3, 463.2, 320.1, 204.1, 191.1, 186.1, 168.1, 138.1, 126.1 |
|  | 656.7863 (+2) | C52H87N11O28 | 1311.557 | 0.46 | 1091.5, 906.4, 763.4, 719.3, 648.3, 534.2, 463.2, 320.1, 204.1, 186.1, 168.1, 138.1, 126.1 |
| 8.11 | 470.7034 (+2) | C37H63N7O21 | 939.3921 | -0.96 | 719.3, 648.3, 534.2, 445.2, 391.2, 302.1, 262.1, 204.1, 186.1, 168.1, 138.1, 126.1 |
|  | 692.3044 (+2) | C55H92N12O29 | 1382.594 | -0.14 | 1162.5, 1073.5, 977.4, 888.4, 834.4, 581.8, 533.3, 302.1, 204.1, 186.1, 168.1, 138.1, 126.1 |
| 10.81 | 497.2168 (+2) | C40H66N8O21 | 992.4186 | -0.7 | 790.3, 701.3, 630.3, 516.2, 462.2, 445.2, 373.2, 302.1, 204.1, 186.1, 168.1, 138.1, 126.1 |
| 12.76 | 895.8787 (+2) | C71H117N13O40 | 1789.736 | 3 | 1551.6, 1366.6, 1348.6, 1223.5, 1163.5, 1020.4, 978.4, 835.4, 204.1, 186.1, 168.1, 138.1, 126.1 |

### Additional structural resolution

*N*AM residues were not reduced prior to LC/MS analysis. This left the potential for mutarotation of their non-reduced glycan ends. However, since MS was used to detect these ions, this step is not necessary. In fact, it is possible that an additional sample preparation step, especially a relatively non-specific acid-base reaction, would have unintended consequences and confound or limit the specificity of the data acquired. This is, of course, at the expense of sensitivity since potentially the signal for one molecular species is divided amongst several chromatographic peaks. An **extracted ion chromatogram** (**EIC**) for *m/z* 895.5-896.5 in one region of the mass chromatogram was used to deduce the structure for another muropeptide (**Fig. 3**). This prediction was determined in the same way as the first structure (**Fig. 2B**), using both the accurate single stage *m/z* measurement as well as the tandem mass spectra from both *Experiments 1* and *2* to add confidence to the assignment. Since all of the measured *m/z* values were within ± 5 ppm of the calculated theoretical *m/z*, and the tandem mass spectra acquired for these three peaks were identical, the same primary structure assignment was made for all three (Note: the position of the non-reduced *N*AM ends are arbitrary in the displayed structure, since these cannot be determined using the data acquired in these experiments).

**Figure 3.**
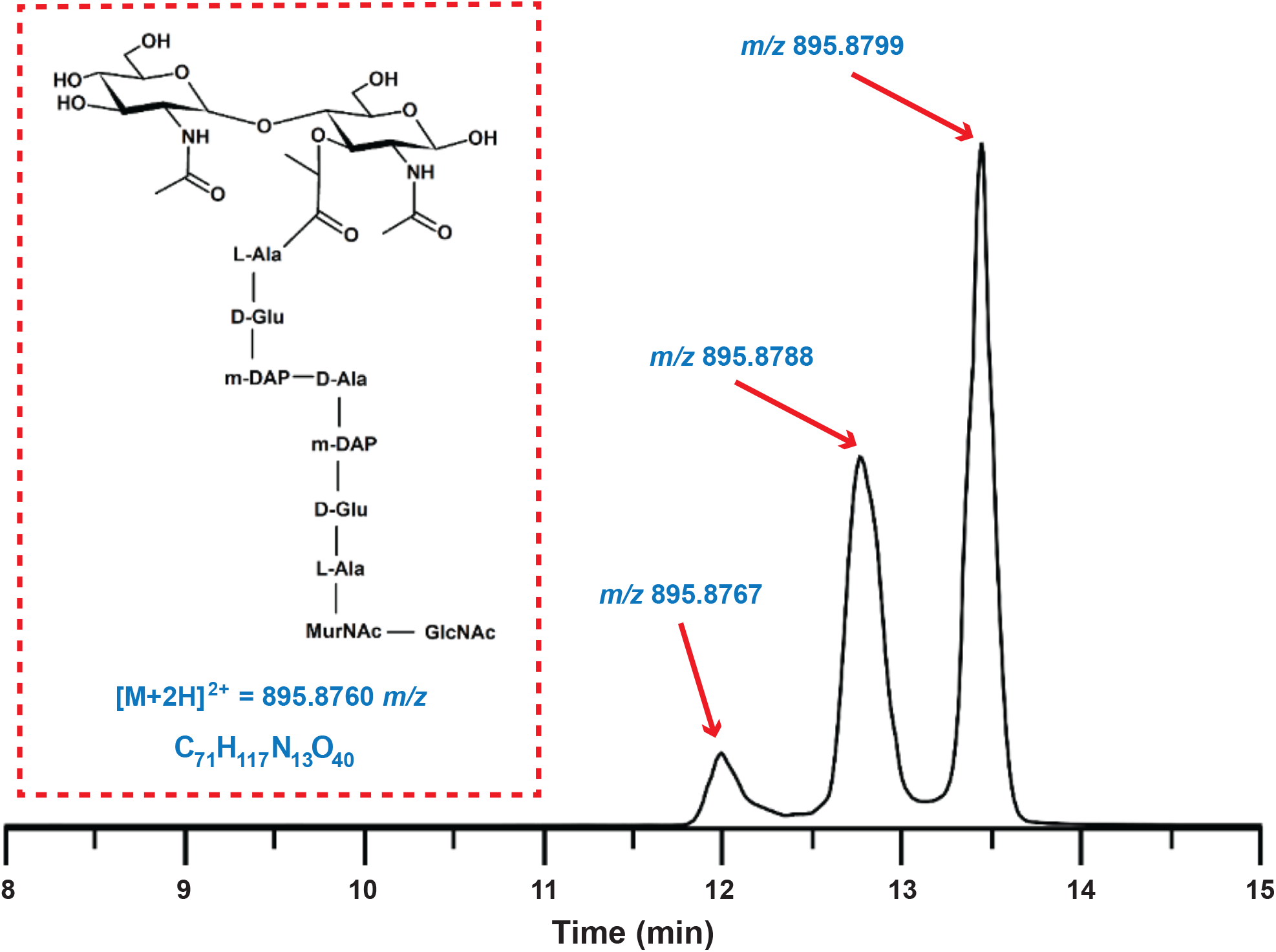
Structure of a crosslinked *R. typhi* PGN subunit. Extracted ion chromatogram of m/z 895.5-896.5 in LC/MS *Experiment 2*. Structure assignment is shown with exact mass for the empirical formula. Three chromatographic peaks were observed, all measured within ± 5 ppm of the calculated mass. These peaks are probably due to mutarotation of the non-reduced glycan ends of *N*AM residues.

### Implications for *Rickettsia* biology

Our results corroborate pathway predictions for *Rickettsia* PGN metabolism (**Fig. 4**) and present the first PGN structure determination for any species of *Rickettsia*. Before now, limited insight on *Rickettsia* PGN was gained from bioinformatics (Gillespie et al., 2009; Driscoll et al., 2017), electron microscopy (Silverman and Wisseman, 1978), and biochemical analysis of individual amino acids within PGN (Pang and Winkler, 1994) from various *Rickettsia* species. More recently, *R. canadensis* was shown to 1) induce NOD1 activation during host cell infection (indicating the presence of meso-DAP), 2) form regular, rod-shaped cells relative to other Rickettsiales species via EM analysis, and 3) incorporate into its cell envelope a d-alanine analog detected with copper-catalyzed click chemistry and microscopy (Atwal et al., 2021). These studies, in conjunction with our work reported here, leave little doubt that *Rickettsia* species synthesize a canonical murein layer. Furthermore, our recent report characterizing *Rickettsia* lipid A structure (Guillotte et al., 2021), coupled with a prior study illustrating the importance for *N*-acetylquinovosamine synthesis in forming the *Rickettsia* O-antigen (Kim et al., 2019), collectively indicate Rickettsiae elaborate a typical Gram-negative bacterial cell envelope that must derive from *N*AG-1-P scavenged from amino sugar metabolism within the host cell cytosol (Driscoll et al., 2017) (**Fig. 4**).

**Figure 4.**
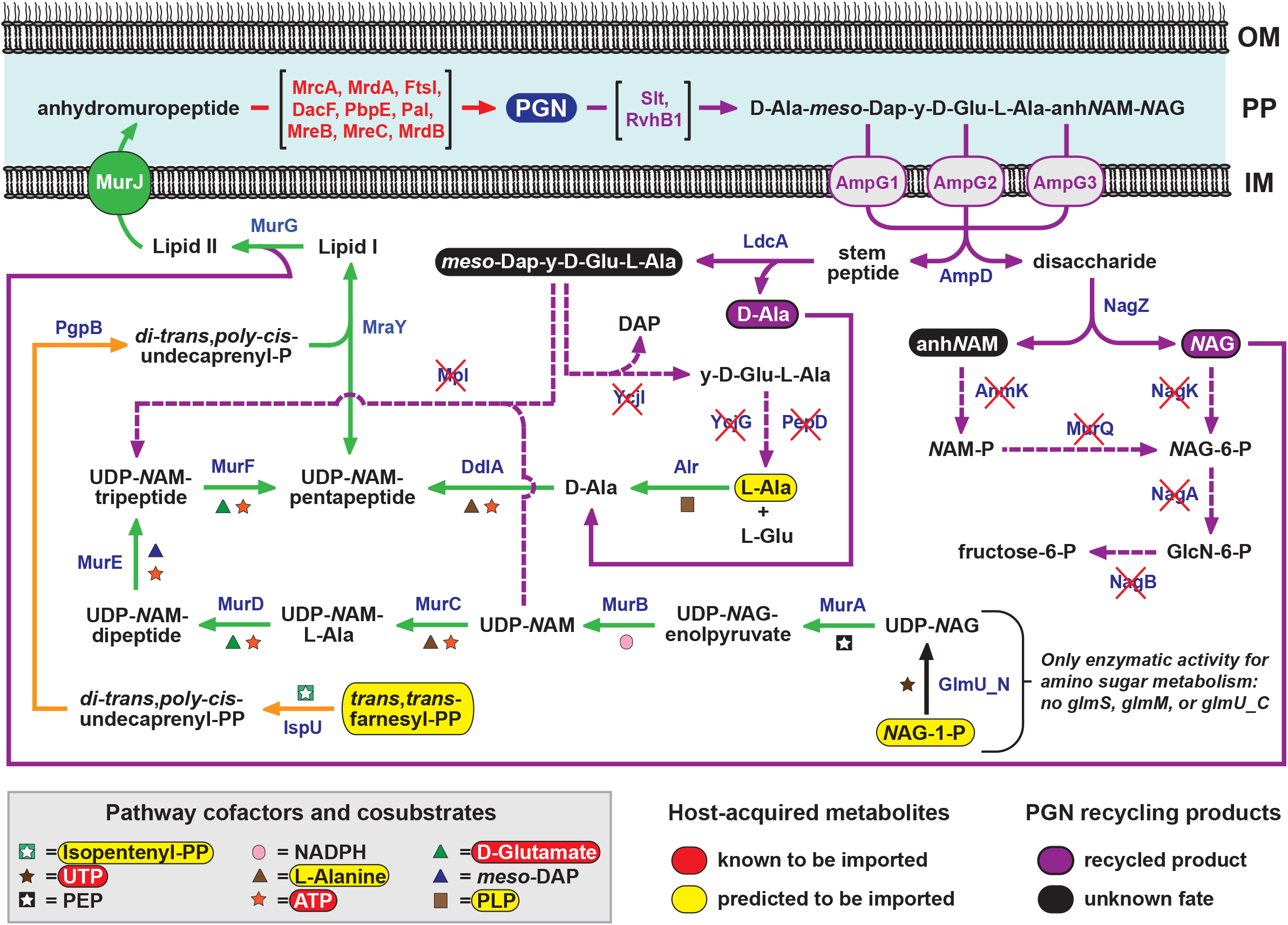
Predicted pathway for *Rickettsia* PGN metabolism. PGN biosynthesis by species of *Rickettsia* requires import of *N*AG-1-P, L-alanine (L-Ala), trans,trans-farnesyl-PP (FPP), and several cofactors from the host cytosol (Driscoll et al., 2017). *N*AG-1-P is converted to UDP-*N*AG) via the uridyltransferase GlmU_N, with the subsequent pathway to Lipid II (green arrows) highly conserved in *Rickettsia* genomes. Via alanine racemase (Alr), imported L-Ala is converted to D-Ala, which is incorporated into the stem peptide by D-alanine--D-alanine ligase A (DdlA). Imported FPP is converted via IspU and PgpB in to di-trans,poly-cis-undecaprenyl-P (orange arrows), which serves as the lipid carrier for Lipid I and Lipid II. At least nine conserved enzymes participate in incorporation of PGN into the murine sacculus (red) (Atwal et al., 2021). The pathway for PGN recycling (purple arrows) initiates with subunit excision via lytic transglycosylases (Slt and RvhB1). Individual subunits are imported back to the cytoplasm via AmpG transporters, which are present in multiple divergent copies in each *Rickettsia* genome (Gillespie et al., 2012a). The full Gram-negative bacterial PGN recycling pathway is shown (Park and Uehara, 2008), with enzymes lacking in *Rickettsia* genomes marked with red Xs. This minimal set of enzymes generates two recyclable products (purple ellipses), while the fate of the remaining degraded products (black ellipses) is unknown.

While our work is a major step forward in understanding a critical component of the *Rickettsia* cell envelope, several questions remain unanswered. It is anticipated that illuminating these unknowns will ultimately guide research on targeting the rickettsial cell envelope with more prudent therapeutics to combat fatal rickettsioses. First, while a recent study suggests host isoprenoids (utilized by MraY to yield Lipid I, see **Fig. 4**) are imported by *R. parkeri* during infection (Ahyong et al., 2019), the nature of isoprenoid (or *N*AG-1-P) uptake by rickettsiae remains unknown. These transport pathways must be determined as they are presumably unique to rickettsiae and should be excellent drug targets. Second, host detection of and insult to PGN during *Rickettsia* infection remains unknown. While our determined structure of *R. typhi* PGN indicates the canonical *N*AG-*N*AM backbone with typical muropeptide crosslinks, the Ala residues observed elongating the muropeptide stems may affect the ability of host detection (NOD1, PGRPs) and/or targeting of PGN via lysozyme and other antimicrobial peptides. Finally, and related to host defense, rickettsiae curiously contain only a minimal part of the well-characterized Gram-negative PGN recycling/turnover pathway, though with amplification of genes encoding the anhydromuropeptide permease AmpG (**Fig. 4**). If functional, PGN turnover will limit host NOD1 detection and surveillance by PGRPs. Also, recycling of at least *NAG* and D-Ala may pose less metabolic burden to infected host cells, considering that the majority of pilfered *N*AG-1-P shunts to LPS biosynthesis (Guillotte et al., 2021), which is trafficked retrograde to the bacterial outer membrane. As bacterial PGN recycling limits the effectiveness of certain cell wall-targeting antibiotics (Mayer et al., 2019), understanding the specific mechanisms employed by different microbes for exoskeleton turnover should lead to improved therapeutics.

## Conclusion

Structures of muropeptides from *R. typhi* PGN were inferred from mass spectra acquired with accurate mass measurements. The basic structure of *R. typhi* PGN was determined to be similar to that of other Gram-negative bacteria, containing mostly a tetrapeptide stem with mDAP residues included. Deviations in stem peptide elongation are worthy of future investigation. This was the first demonstration of PGN structure analysis for *R. typhi*, and indeed, the first PGN structure determined for any *Rickettsia* species. Rickettsiae are difficult organisms to culture because of their obligate intracellular life cycle, and isolation of compounds from them is complicated by their co-culture with host cells. Since PGN has been a drug target for many years and its structure affects the efficacy of PGN-targeting drugs, chemical structure analysis of PGN is paramount for successful drug discovery efforts. Additionally, the study of basic biology of Rickettsiae will benefit from this research by increasing interest in host metabolite thievery and combining it with efforts to develop a more complete picture of obligate intracellular parasitism.

## Acknowledgments

We are grateful to Dr. Jeanne Salje for discussions on peptidoglycan biology. DRG thanks the University of Victoria-Genome BC Proteomics Centre and is grateful to Genome Canada and Genome British Columbia for financial support for Genomics Technology Platforms (GTP) funding for operations and technology development (264PRO) as well as NIH’s NIAID for funding 1R01 AI147314-01A1. This work was also supported with funds from NIAID grants R01AI017828 and R01AI126853 (AFA), R21AI26108 and R21AI146773 (JJG and MSR), and R21AI156762 (JJG). KER-B was supported in part by NIH/NIAID grant T32AI095190 (Signaling Pathways in Innate Immunity). The content is solely the responsibility of the authors and does not necessarily represent the official views of the funding agencies. The funders had no role in study design, data collection and analysis, decision to publish, or preparation of the manuscript.

